# Proximity map for p53 unveils a strong link with PML Nuclear Bodies in HEK293 cells

**DOI:** 10.1101/2025.10.22.683840

**Authors:** E. De Bruycker, G. Vandemoortele, D. De Sutter, GD. Moschonas, S. Eyckerman

## Abstract

The tumor suppressor p53 is among the most studied proteins in cancer, with mutations in nearly half of all tumors. Even when *TP53* is intact, its regulatory pathways are often disrupted, underscoring its central role in tumorigenesis. To explore how cellular context shapes the subcellular environment of p53, we used BioID proximity labeling in HEK293 cells. These experiments revealed a striking enrichment of proteins involved in SUMOylation and proteins that reside in PML nuclear bodies. A strong enrichment of the adenoviral E1B protein, originally used for transformation of this cell line, may explain this striking observation, and can shed light on the remarkable difference with the proximity map of p53 in the HCT116 colon carcinoma cell line. Strong enrichment of the mediator complex in HCT116, related to active p53-dependent transcription, is missing in 293 cells and can explain the lack of p53 transcriptional activity in these cells. This study emphasizes how cellular transformation can affect the proximal proteome of a key protein in cell cycle control, DNA repair, apoptosis, senescence and metabolism, and confirms previously reported inhibition of p53-dependent transcription by sequestering the protein in PML nuclear bodies.

## Introduction

Tumor suppressor protein p53 is arguably the most extensively described protein in biomedical literature. Often referred to as ‘the guardian of the genome’, it is the most frequently mutated protein in the context of tumorigenesis (in approx. 50% of the cases). In case the *TP53* gene remains intact, pathways controlling the activity of the protein are almost invariably found to contain mutations (e.g. amplification of the inhibitory *MDM2* gene)^1^, demonstrating the critical role of p53 in cancer.^2^ Despite extensive research, the study of p53 continues to evolve, with new findings regularly contributing to the already considerable data pool^3^. p53 is in essence a transcriptional activator regulating the expression of genes involved in cell cycle progression, DNA damage repair, apoptosis, cellular senescence and non-canonical activities (e.g. metabolism and autophagy).^4, 5^ In a number of specific cases, such as in response to severe DNA damage, p53 has been shown to localize towards the mitochondria, where it interacts with pro-apoptotic proteins. This association triggers the release of cytochrome C and activation of caspases, ultimately leading to apoptotic cell death.^6-8^ Due to the central role of p53 in the cell, its activity is strongly regulated by post-translational modifications (PTMs), such as acetylation and phosphorylation, ubiquitination and SUMOylation.^9^ In non-stressed cells, the intracellular levels of p53 are tightly regulated by its interaction with E3 ubiquitin ligases, primarily Mdm2, which targets p53 for proteasomal degradation. Upon pro-oncogenic events such as genotoxic stress, temporary stabilization of p53 is promoted through acetylation and phosphorylation by acetyltransferases (e.g. CBP/p300 complex) and kinases (e.g. ATM, ATR, Chk2 and HIPK2), respectively.^10^

Several reports reveal a cell-specific role for Promyelocytic Leukemia Nuclear Bodies (PML NBs) in the regulation of p53. PML NBs are membrane-less organelles localized in the nucleus of all mammalian cells, formed through liquid-liquid phase separation (LLPS).^11, 12^ They harbor proteins involved in crucial cellular processes such as transcriptional regulation, DNA damage repair, apoptosis, anti-viral response and tumor suppression.^13, 14^ PML NBs are assembled upon SUMOylation of their essential scaffold protein, PML.^15^ Small Ubiquitin-like Modifier (SUMO) modifications involve small ubiquitin-like proteins that are conjugated to lysine ε-amino groups accessible on the surface of cellular proteins. The covalent attachment of a SUMO group is orchestrated by an enzymatic cascade involving the E1 enzyme (UBA2) activating SUMO, the E2 conjugating enzyme Ubc9 and is further regulated by E3-like factors.^16^ This PTM serves as a docking site for proteins harboring SUMO interacting motifs (SIMs).^17^ Upon stress stimuli such as viral infection, oxidative or oncogenic stress, target proteins undergo SUMOylation, resulting in their recruitment to the core of PML NBs through SUMO-SIM interactions. In response to DNA damage, p53 is SUMOylated resulting in its translocation towards PML NBs where p53 undergoes additional modifications and becomes activated by transiently PML NBs-localized client proteins such as HIPK2, ATR and CBP/p300.^13^ This p53 response pathway is further regulated by viral proteins. For instance, the oncogenic E1B-55K adenoviral protein sequesters and inhibits p53 in PML-NBs.^13^

The intrinsic disadvantage of cell lysis for studying protein-protein interactions is the unavoidable mixing of proteins from different cellular compartments. Hence, the identification of compartment-specific proteins and/or processes is challenging using cell lysis-dependent proteomic techniques. Proximity-dependent biotinylation by methods such as BioID offers the opportunity to circumvent this hurdle through tagging proteins prior to cell lysis.^18^ BioID depends on a constitutively active, *E. coli*-derived biotin ligase (BirA*) fused to the bait protein. Upon biotin administration, proximal proteins within an approximate range of 10 nm will become biotinylated. This labelling facilitates efficient isolation and purification via streptavidin afterwards. Furthermore, this approach avoids the inherent limitations of immunoprecipitation and affinity purification strategies, such as the disruption of transient and indirect interaction partners.^19^ BioID has emerged as an invaluable tool for characterizing known membrane-less organelles including stress granules and the nucleolus^20-22^, and for revealing new biomolecular condensates^23^.

In this study we have applied BioID proximal labelling to identify vicinal proteins of p53 in HEK293 cells. After treatment of the cells with the chemotherapeutic agent doxorubicin, we observed strong associations with proteins involved in p53’s canonical transcriptional role. In addition, a strong link with SUMOylation and PML NBs could be observed. However, these factors are not detected in the endogenous p53 proximal proteome of spontaneously immortalized HCT116 cancer cell line. Both the SUMO link and association with PML NBs are likely explained by the presence of the adenoviral oncogenic E1B protein interacting with p53 in HEK293 cells.

## Results

### Proximal labelling of p53 with BioID in HEK293 cells confirms known interactions

The essential role of p53 in numerous biological processes, its frequent alteration in human cancers and its extensive characterization through interactome studies using classical methods in different cellular systems, inspired us to revisit the interactome of p53 using BioID.^24, 25^ As a model system, we used HEK293, one of the most commonly used cell lines in cell biology, which is characterized by a wild-type p53 status unlike many other cell lines.^26^ To fine-tune the expression level of p53-BirA* to match endogenous levels, the doxycycline inducible FlpIn T-REx cell line was employed, regularly used in BioID studies.^20, 27-29^ As a control set-up, we generated a cell line expressing BirA* fused to a nuclear localization signal (NLS), enabling labeling of nuclear background proteins. This setup allows for elimination of non-specific background nuclear proteins from the p53 proximal proteome (Figure 1A).

**Figure 1.**
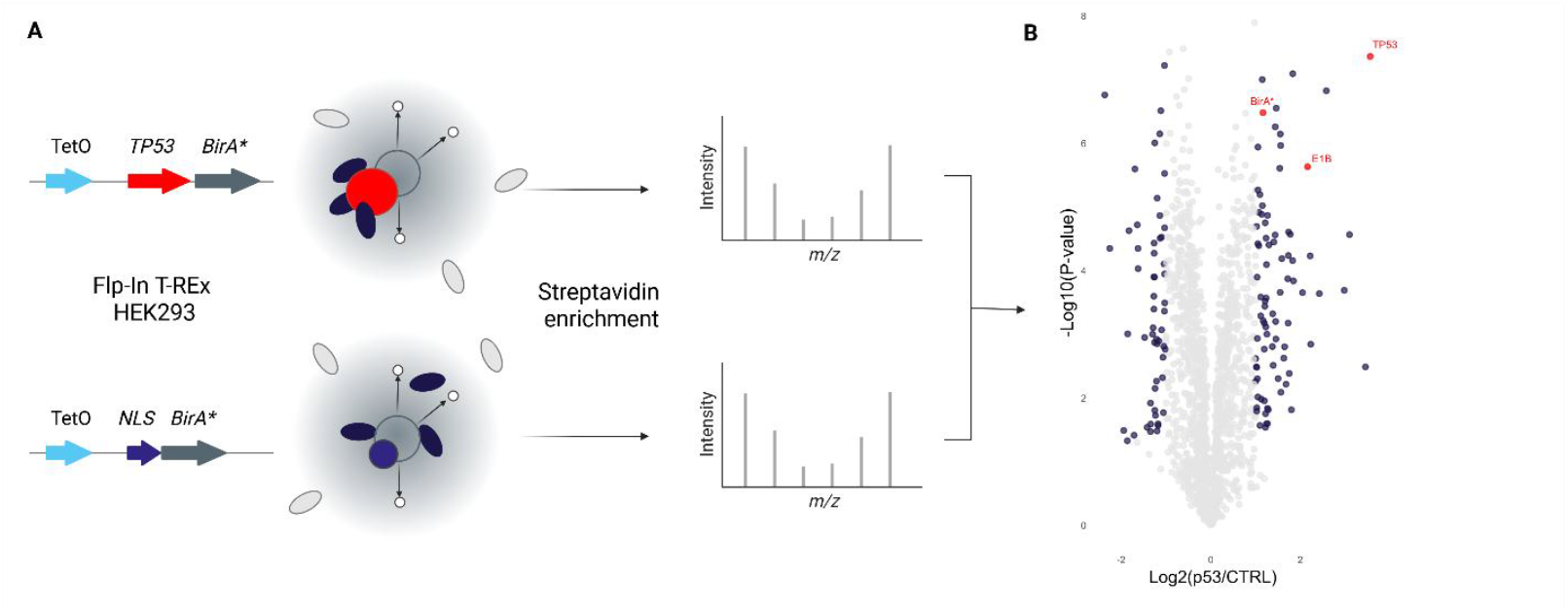
BioID for p53-BirA* in HEK293 cells. (A) Experiment overview. Generation of Flp-In T-REx 293 cells expressing p53-BirA* and NLS-BirA* under a doxycycline-inducible promoter (TetO). This allows for proximal proteome labelling and precipitation using streptavidin beads. Enriched proteins are quantified through mass spectrometry-based proteomics. (B) Volcano plot depicting p53-BirA* (right) versus NLS-BirA* (left). All proteins with p-value ≤ 0.05 and |log2(FC)| > 1 are indicated in blue. In red, bait proteins p53 and BirA*, and E1B.

After doxycycline induction of the bait protein, and upon activation of the DNA damage response with doxorubicin, proximal labelling was performed by treating the cells with biotin followed by enrichment and analysis through mass spectrometry. A total of 2,129 Protein Groups were detected of which 1,566 were quantified. A total of 86 proteins were significantly enriched for p53-BirA* compared to control (p-val ≤ 0.05, Log2FC>1; Figure 1B, Supplementary Table 1). High enrichment of BirA* in the p53-BirA* vs. the NLS-BirA* condition is explained by strong p53 labeling while BirA* autobiotinylation is absent or low^30^. Furthermore, 51 out of 86 (60%) significantly enriched proteins have been demonstrated to interact with p53 according to the BioGRID^4.4^ database (Figure 2A, Supplementary Figure S1). This underscores the reliability of BioID in providing a relevant view on the proximal interaction network of p53.

**Figure 2.**
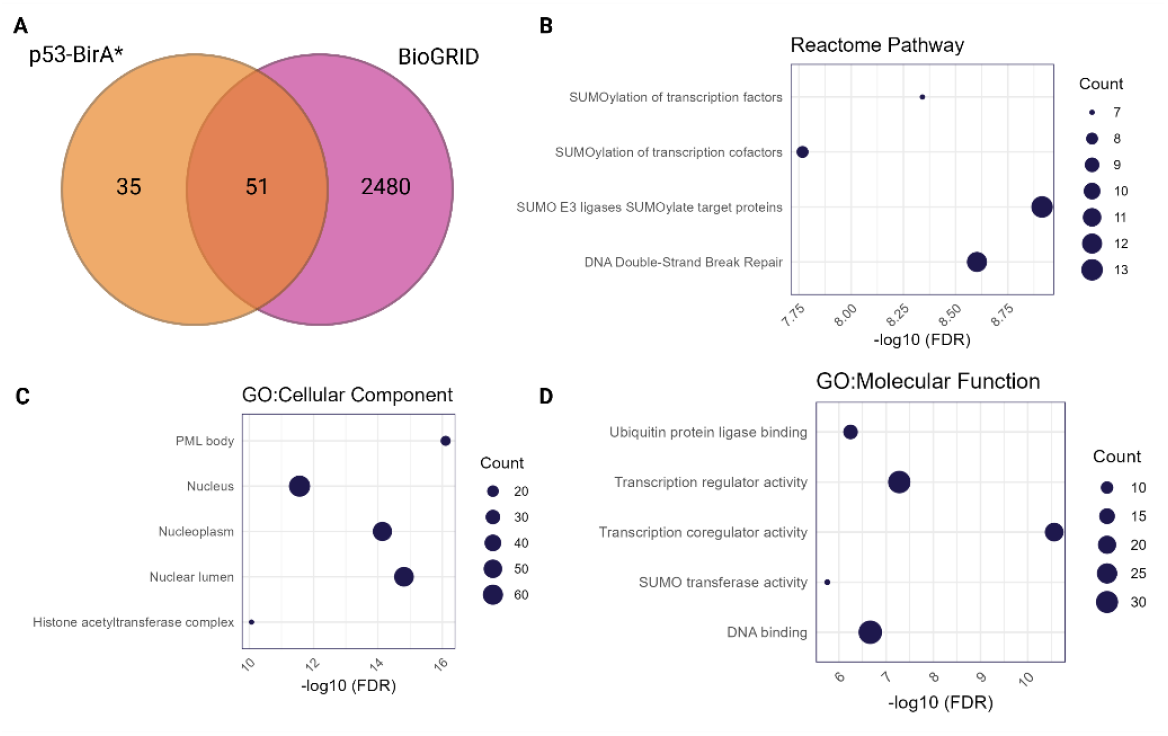
Overlap with BioGRID and Gene Ontology Enrichment analysis for differentially enriched proteins for p53-BirA*. (**A**) Overlap with BioGRID^4.4^ repository. Majority of the enriched proteins for p53-BirA* (60%) have been described. (**B**) Enriched Reactome pathways. Pathways associated with SUMOylation and DNA repair are enriched. (**C**) Cellular Component demonstrates association with PML (nuclear) body. (**D**) Molecular Function. Enriched proteins for p53 are involved in transcription.

As the HEK293 cell line was originally established through stable integration of adenoviral DNA encoding for E1A an E1B proteins^31^, and as E1B has been demonstrated to interact with p53^32, 33^, we have added these proteins to the protein search list used for MS-based peptide identification. Not surprisingly, we detected E1B, but not E1A, as one of the most significantly enriched proteins for p53 (Figure 1B).

### Proximal labelling of p53 in HEK293 reveals a link with PML nuclear bodies

To further explore the results, a gene ontology enrichment analysis was performed using the STRING-Database (version12.0) on the full set of 86 proteins. Enriched terms for Reactome pathway, Cellular Component and Molecular Function are illustrated in Figure 2. Reactome pathway analysis suggests that the proteins enriched for p53 are functionally involved in DNA repair mechanisms, as indicated by ‘DNA Double-Strand Break Repair’ (2.50E-9). This pathway coincides with the effects of doxorubicin treatment and represents one of the major pathways p53 is involved in.^34^ Relevant proteins for this pathway are ERCC1 and ERCC4, POLH, RAD50, BLM and TP53BP1 (Figure 3A). Furthermore, the proximal partners of p53 in HEK293 cells show a strong link to ‘SUMOylation’ as indicated by highly significant FDR scores for ‘SUMO E3 ligases SUMOylates target proteins’ (1.22E-9) and ‘SUMOylation of transcription factors and cofactors’ (4.56E-9; 1.71E-8). Proteins enriched in the SUMO pathway are TOPORS, PIAS1, -2 and -4, and SUMO1 (Figure 3A). Gene Ontology enrichment for Molecular Function agrees with the SUMO link as illustrated by the ‘SUMO transferase activity’ (1.75E-6) GO term encompassing the proteins PIAS1, -2 and -4, TOPORS and SUMO1. Furthermore, ‘Ubiquitin protein ligase binding’ (5.65E-7) may be enriched because of the SUMO link, since SUMO is a small Ubiquitin-like Modifier protein. Other terms are associated with the major role of p53 as transcription factor: ‘Transcription coregulator and regulator activity’ (2.75E-11; 5.24E-8) and DNA binding (2.16E-7).

**Figure 3.**
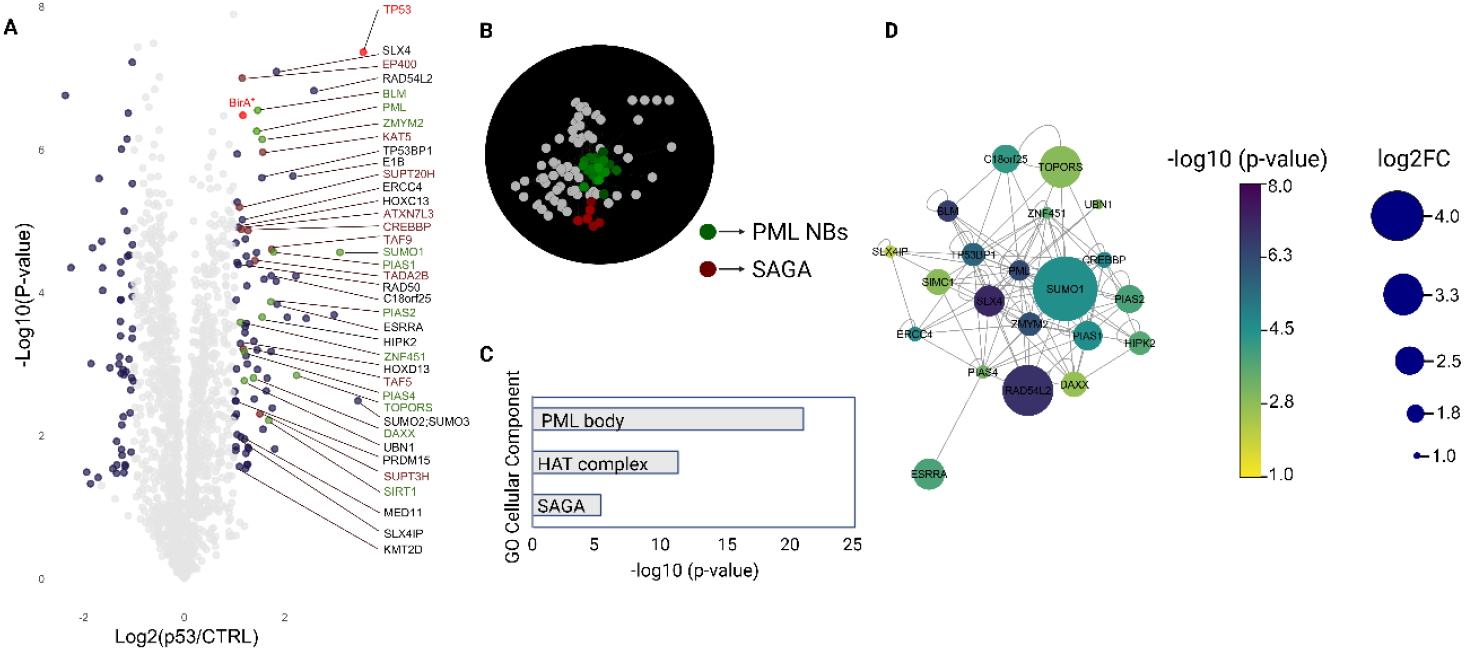
BioID demonstrates a link with PML nuclear bodies in HEK293 cells. (**A**) Volcano plot illustrating the enriched proteins for PML Nuclear Bodies and SAGA complex. Proteins are colored according to the SAFE analysis. (**B**) SAFE analysis demonstrating the major biological networks in the dataset for p53-BirA*. (**C**) Proteins corresponding to PML nuclear body by SAFE analysis. Each protein in the network is colored according to the -log10 (p-value) and log2 (fold change) in the volcano plot.

In addition, using g:Profiler^35^ and the spatial analysis for functional enrichment (SAFE) tool in cytoscape^36^, biological networks were visualized for the enriched proteins for p53 (Figure 3B). As already hinted by Cellular Component analysis, the major network enriched for p53-BirA* is ‘PML (nuclear) body’, indicating that p53 resides in these nuclear bodies in HEK293 cells upon DNA damage. A total of 21 proteins were found to be associated with PML nuclear bodies, including the core component protein PML (Figure 3A, B). Furthermore, the Spt-Ada-Gcn5 acetylatransferase (SAGA) complex, involved in gene regulation, chromatin modification and DNA damage repair, is the second enriched complex (Figure 3A, B).^37, 38^

### p53 associates with PML nuclear bodies in HEK293 but not in HCT116 cells

The strong enrichment of terms related to SUMOylation in combination with the association with PML nuclear bodies directed our search further. We revisited a BioID experiment performed in the HCT116 cancer cell line, having a wild-type p53 status^26^, previously published by our research group (PXD011702). In this study, the BirA* module was introduced on the endogenous protein by CRISPR-Cas9 engineering. An elegant system based on the T2A autocleavage peptide was used as a control set-up to remove background proteins from the analysis (‘T2A-Switch’ design).^39^ Using the same cut-off values (p-value < 0.05, |log2FC| > 1), a total of 78 proteins were enriched for p53 compared to the control setting in the HCT116 cell line (Figure 4A). In this dataset, the enriched biological networks correspond to the Mediator Complex and SAGA complex. Cellular Component analysis reveals that the proteins are primarily involved in histone acetyltransferase (HAT) and again Mediator and SAGA complex.

**Figure 4.**
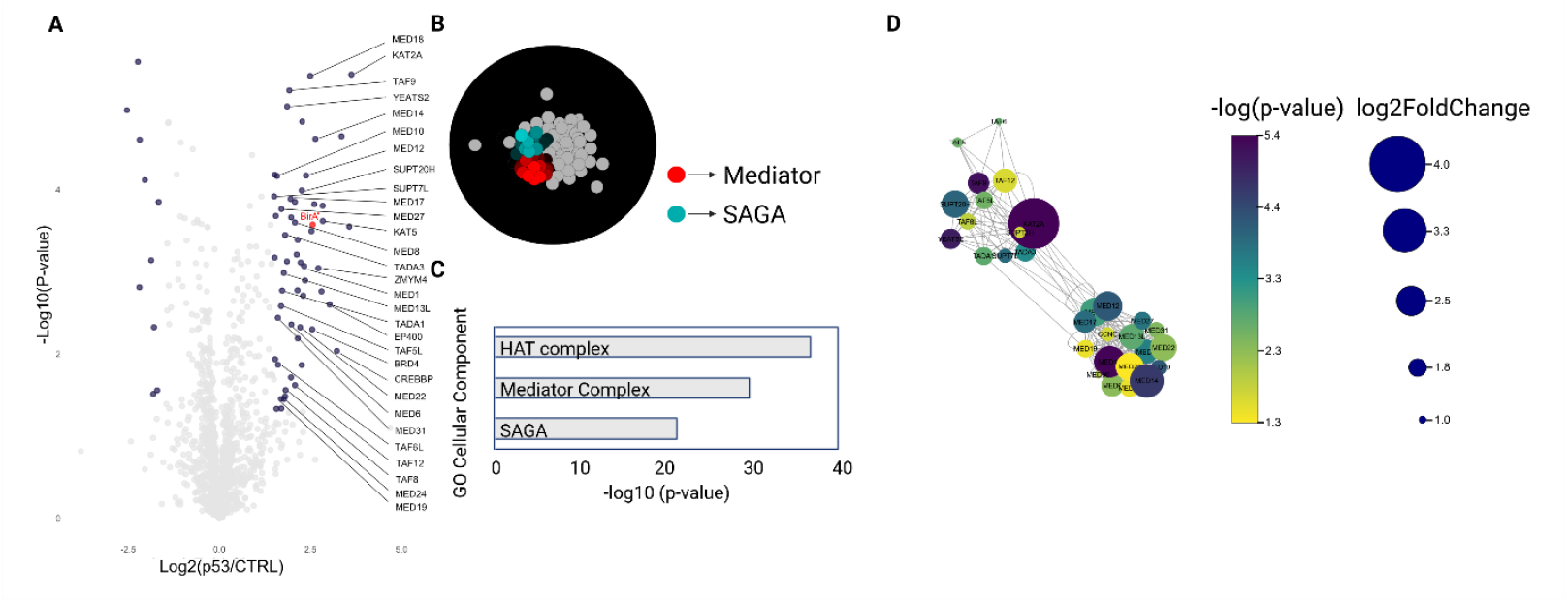
Re-analysis of published BioID data for p53 in HCT116 cell line. (**A**) Volcano plot. All significantly enriched proteins (p-value < 0.05, |log2FC| > 1) are indicated in blue. (**B**) SAFE analysis. Mediator complex and SAGA complex are among the strongest enriched biological networks. (**C**) Cellular Component analysis. (**D**) Mediator complex and SAGA complex according to SAFE analysis. Color and size of the proteins represent the p-values and fold change in the volcano plot.

However, no link with ‘SUMO’ nor PML nuclear bodies could be observed in HCT116 for p53. Therefore, both datasets were compared considering all significantly enriched proteins (FDR=0.01). A total of 62 proteins overlapped for both datasets largely correspond to the HAT complex, illustrating that this interaction occurs in both cell lines. Surprisingly, proteins associated with PML nuclear body were exclusively enriched in the HEK293 cell line (Figure 5).

**Figure 5.**
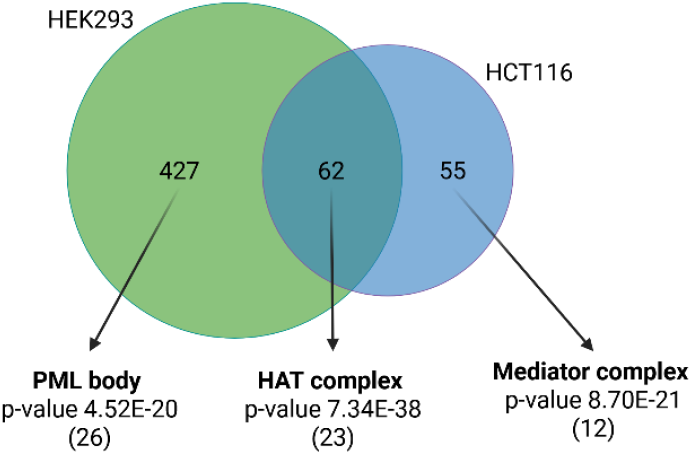
Venn diagram illustrating the overlap of significantly enriched proteins in the HEK293 and HCT116 dataset for p53-BirA*. With an FDR of 0.01, 489 and 117 proteins are enriched for p53 in the HEK293 and HCT116 dataset, respectively. A total of 62 proteins overlap. The strongest GO term associated with this subset is the HAT complex. For the HEK293 cell line, 427 proteins are exclusively enriched. A total of 26 proteins are associated with PML bodies (Figure S1, left panel), and 55 proteins were exclusively enriched in the HCT116 dataset of which 12 (Figure S1, right panel) were associated with the mediator complex.

## Discussion

Intensive research on p53 has resulted in a staggering list of 2,531 protein partners according to the BioGRID repository (version 4.4, ^40^; April 2025). To gain a better understanding of the interaction landscape of p53, we aspired to generate a proximity map in the HEK293 cell line using proximal labelling with BioID. We employed the widely used Flp-In T-REx 293 cell line that allows efficient recombinase-assisted generation of a doxycycline-inducible isogenic population carrying a single integrant in a well characterized locus. To allow elimination of background biotinylation, a control cell line was generated expressing BirA* targeted to the nucleus by a NLS signal peptide. A total of 86 proteins were significantly enriched for p53 (pval≤0.05, |log2FC|>1) compared to control. The majority of the enriched proteins largely overlap with the protein partner list for p53 in the BioGRID repository^4.4^ (60%, Figure 2). Despite this long interactome list, a set of novel candidate partners (35) was identified. However, the majority of these novel candidates map to protein complexes that have been described before to associate with p53. These results thus expand the p53 interactome to include interactions that are likely indirect, a known feature associated with BioID proximal labeling.

The proximity map obtained under conditions of genotoxic stress (i.e. doxorubicin treatment) revealed the expected transcriptional (co-) regulator complexes and histone acetyltransferase (HAT) complex. The HAT complex plays an important role in regulating the p53 pathways, including cell cycle, DNA damage response, apoptosis and metabolism.^41^ Among the highly enriched proteins, Histone/Lysine acetyltransferase KAT5, commonly referred to as Tip60, plays a key role in transcriptional activation by acetylating lysine residues on histones H4 and H2A.^42^ Beyond its function in gene activation, KAT5/Tip60 has been implicated as a central player in DNA damage repair. ^43^ KAT5/Tip60 and cofactor TRAPP bind chromatin adjacent to DNA double strand breaks in the ATM-dependent DNA damage signaling cascade. Through hyperacetylation of histone H4 by KAT5, DNA repair proteins accumulate in close proximity of the DNA damage site.^44^ Furthermore, Bloom syndrome protein (BLM) was detected. This is a member of the RecQ family of DNA helicases, facilitating repair of DNA double strand breaks. It has been demonstrated that BLM interacts with p53BP1, forming a scaffold that facilitates the recruitment of p53 towards γH2AX-marked sites. ^45^ In addition, structural maintenance of chromosome (SMC) family member (Rad50) and Nibrin (NBN) were significantly enriched. They are members of the MRN (MRE11, Rad50, NBN/Nbs1) complex, which is an essential component in DNA double strand break repair, forming MRN DNA foci at γH2AX marked sites.^46-48^ Lastly, homeodomain-interacting protein kinase 2 (HIPK2) has been reported to monitor DNA damage by targeting p53.^49^

A strong enrichment of proteins involved in SUMOylation could be observed in HEK293, but not HCT116, which can be linked to PML nuclear bodies. Besides their role in protein SUMOylation, PML NBs serve as storage depots for proteins, releasing the proteins upon specific stimuli.^11, 12^ Moreover, p53 is recruited to PML NBs upon DNA damage induced by amongst others doxorubicin.^50^ Once recruited, p53 meets many of its key regulatory enzymes also enriched in our dataset, including HIPK2, KAT5, CREBBP and SIRT1.^51^ Furthermore, our dataset revealed the presence of TOP1 Binding Arginine/Serine Rich Protein (TOPORS) and Protein inhibitor of activated STAT (PIAS) 1, 2 and 4. TOPORS is a E3 SUMO1 protein ligase that SUMOylates p53, which results in a higher stability.^52^ Members of the PIAS family are E3 SUMO ligases known to modify p53. However, the functional consequence of p53 SUMOylation by the PIAS protein family remains controversial, meaning it can induce both inactivation by promoting nuclear export or accumulation by increasing its stability.^53^ In addition, Histone cell cycle regulator (HIRA) is part of the HUCA (HIRA/UBN1/CABIN1/ASF1a) complex that is responsible for histone H3.3 modifications at sites of DNA repair and transcriptional active genes involved in cell cycle control.^54^ The high recruitment affinity of H3.3 regions stimulates the localization of Death-domain associated protein 6 (DAXX), which allows for DAXX-mediated localization of PML NBs towards genomic target sites.^54, 55^ Hence, both HIRA and DAXX trigger PML NBs to localize towards DNA damaged regions and genes involved in cell cycle. ^54, 55^

Oncogenic viruses such as human papilloma virus (HPV), simian virus 40 (SV40) and adenovirus were instrumental in helping to understand the critical role of p53 in preventing oncogenic transformation.^56^ E1B and E1A are oncoproteins expressed by Human Adenovirus 5 that cause the transformation of various cell lines, including HEK293 cells.^57^ They affect the functions of main cell cycle regulators, p53 and pRB, and proteins in the apoptotic pathway, thereby allowing immortalization of cell lines.^58^ E1B SUMOylation regulates its nucleo-cytoplasmic shuttling and intra-nuclear localization to exert the transformation of mammalian cell lines. Furthermore, evidence was also provided that E1B not solely serves as a target of SUMO, but also stimulates SUMO modification of p53 in PML NBs.^59^ This results in an increased stability of p53’s association with PML NBs at first, and subsequently, results in maximal repression of p53’s transcriptional activity through facilitating its nuclear export towards cytoplasmic aggresomes.^13, 60^

This study provides further insights into the p53 proximal interactome and sheds light on p53 regulation, which critically depends on the cellular background. Moreover, our results highlight the critical importance of choosing an appropriate cellular model for interactomics research, especially when performed on proteins that are targeted in the process of cellular transformation, thereby emphasizing the potential impact of the immortalization process on the surrounding protein environment, which could influence interaction dynamics and experimental outcomes.

## Materials & Methods

### Plasmids and cloning procedures

The construct used for stable transfection was generated by gateway cloning of TP53 ORFeome entry clone #3774 to the pDEST-pcDNA5-BirA*-FLAG C-term destination vector. The pcDNA5-HA-SV40 NLS-BirA*-FRT/TO was created by PCR amplification of the BirA* coding sequence from the pDEST-pcDNA5-BirA*-FLAG C-term destination vector with a 5’ primer encoding the SV40 NLS sequence preceded by a HA tag. The amplicon was then inserted in the pcDNA5-FRT/TO vector by In-Fusion® cloning (Takara Bio).

### Cell culture

Flp-In T-REx 293 cells (Catalog No. R78007, Thermo Fisher Scientific) were cultured in DMEM (Catalog No. 61965026, Gibco) supplemented with fetal bovine serum (FBS, Catalog. No. 10270106, Gibco) at a final concentration of 10%, 12,500 units penicillin-streptomycin (Catalog No. 15070063, Gibco) and HEPES (Catalog No. 15630056, Gibco) at a final concentration of 10 mM. Mycoplasma PCR testing was done to confirm absence of mycoplasma (Minerva Biolabs, MIN-11-1100). Flp-In T-REx 293 cells were generated by seeding 2.81×10^5^ cells for calcium phosphate transfection the next day. 32h after transfection with pDEST-pcDNA5-p53-BirA*-FLAG or pcDNA5-FRT-TO-HA-NLS-BirA*-FLAG, cells were transferred to a 100 mm culture dish and cultured in the presence of 15 µg ml^-1^ blasticidin (Invivogen, ant-bl-1) and 50 µg ml^-1^ hygromycin B (Invivogen, ant-hg-1) for two weeks. Two weeks prior to seeding Flp-In T-REx 293 cells for a BioID experiment, growth medium was changed to DMEM medium supplemented with 10% dialyzed fetal bovine serum (Catalog No. 26400044, Gibco) to remove excess biotin from the medium.

### Proximal labelling with BioID

Three 150 mm culture dishes (Catalog No. 249964, Nunc) were seeded with 6.75×10^6^ Flp-In T-REx 293 pDEST-pcDNA5-p53-BirA*-FLAG cells or with Flp-In T-REx 293 pcDNA5-FRT-TO-HA-NLS-BirA*-FLAG cells for every sample. Three replicates were prepared for each condition. The next day, cells were treated with a low dose of doxycycline (3 ng ml^-1^) to induce the expression of the p53-BirA* fusion and the NLS-BirA*. After 24h, 1 µM doxorubicin (Catalog No. D1515, Sigma) was added to evoke the DNA damage response and induce p53 expression, and 50 µM biotin (Catalog No. B4639, Sigma) was added at the same time to allow biotinylation. The complemented growth medium was replaced with standard growth medium after another 24h to remove excess of free biotin. Three hours later, cells were washed with ice-cold Phosphate Buffered Saline (PBS) (Catalog No. 14190094, Gibco) before detachment in PBS using a cell scraper. BioID purification was performed as described earlier^39^. In brief, cell lysis occurred in 5 ml of ice-cold RIPA lysis buffer containing 250 U of benzonase (Sigma, E1014) by incubation with agitation for 1h on 4°C before sonication (30% amplitude, 5 x 6 s burst, 2 s interruption). The total protein concentration was determined by the Bradford protein assay to normalize input material. For every sample, 90 µl pre-washed Streptavidin Sepharose High Performance bead suspension (Catalog No. 17-5113-01, GE Healthcare) was used for enrichment of biotinylated proteins using a rotator at 4°C for 3h. Afterwards, beads were washed extensively and purified material was resuspended in 20 µl 20 mM Tris-HCl pH 8.0 before addition of 1 µg trypsin (Catalog No. V5111, Promega). After overnight digestion, supernatant was transferred to a mass spectrometry vial and an additional 500 ng trypsin was added for 3h before acidification. All experiments were performed in biological triplicate for downstream LFQ analysis.

### LC-MS/MS instrument and analysis settings

Peptide mixtures were analyzed by LC−MS/MS on an Ultimate 3000 RSLC nano LC (Thermo Fisher Scientific) in-line connected to a Q-Exactive mass spectrometer (Thermo Fisher Scientific). The peptides were first loaded on a trapping column (made in-house, 100 μm internal diameter (I.D.) × 20 mm, 5 μm beads C18 Reprosil-HD, Dr. Maisch, Ammerbuch-Entringen, Germany). After flushing the trapping column, peptides were loaded in solvent A (0.1% formic acid) on a reverse-phase column (made in-house, 75 µm I.D. x 250 mm, 3 µm Reprosil-Pur-basic-C18-HD beads packed in the needle, Dr. Maisch, Ammerbuch-Entringen, Germany) and eluted by an increasing concentration solvent B (0.1% formic acid in acetonitrile) using a linear gradient from 2% solvent B up to 55% solvent B in 120 min, followed by a washing step with 99% solvent B, all at a constant flow rate of 300 nl min^-1^. The mass spectrometer was operated in data-dependent (DDA), positive ionization mode, automatically switching between MS and MS/MS acquisition for the 5 most abundant peaks in a given MS spectrum. Source voltage was set at 3.4 kV, with a capillary temperature of 275°C. One MS1 scan (m/z 400−2,000, AGC target 3×106 ions, maximum ion injection time 80 ms), acquired at a resolution of 70,000 (at 200 m/z), was followed by up to 5 tandem MS scans (resolution 17500 at 200 m/z) of the most intense ions fulfilling predefined selection criteria (AGC target 5 × 104 ions, maximum ion injection time 80 ms, isolation window 2 Da, fixed first mass 140 m/z, spectrum data type: centroid, underfill ratio 2%, intensity threshold 1.3E4, exclusion of unassigned, 1, 5-8 and >8 positively charged precursors, peptide match preferred, exclude isotopes on, dynamic exclusion time 12 s). HCD collision energy was set to 25% normalized collision energy and the polydimethylcyclosiloxane background ion at 445.120025 Da was used for internal calibration (lock mass).

### Mass Spectrometry Data Processing and Interpretation

Obtained raw files (.raw) were analyzed using MaxQuant. Spectra were searched against the human UniProt sequence database. Additional FASTA files for E1B (P03244), E1A (P03254), BirA (ID Z00001) and BirA* (ID Z00002) sequences were included in the searches. Methionine oxidation, N-terminal acetylation, and lysine biotinylation were set as variable modifications, with a maximum of five modifications per peptide. The minimum peptide length was set at seven amino acids, and a maximum peptide mass of 4,600 Da was used. PSM, protein, and site false discovery rates were set at 0.01. The minimum label-free quantitation (LFQ) ratio count was 2, and the Fast LFQ option was disabled. Mass accuracies of 20 and 4.5 ppm were used for the first and main searches, respectively. After the completion of searches, LFQ intensities were loaded in Perseus (version 1.6.2.2) for further analysis.

Proteins only identified by site, reverse hits, and Perseus contaminant proteins were removed from the data matrix. Samples were annotated on condition of analysis. Retained LFQ intensities were transformed to log2 scale, and two valid values were needed in order for the protein to be retained for further analysis. Missing values were imputed from a normal distribution of intensities (0.3 width, 1.8 downshift). Samples were normalized based on Z-scores. A two-sided *t* test (0.01 FDR, 250 randomizations) was performed to reveal differentially enriched proteins in the volcano plot.

### String Database

Significantly enriched proteins were uploaded in the STRING Database (version 12.0, stringr-db.org/) and searched against known human interactions. A full STRING network was generated with the edges indicating both functional and physical associations. Active interaction sources applied were Experiments, Databases, Gene Fusion and Co-Occurrence. The minimum required interaction score was set at medium confidence (0.400). Top GO terms for cellular component and molecular function, and REACTOME pathways are listed in Figure 2.

### p53 SAFE network

SAFE network analysis was performed for the significantly enriched p53-proximal proteins (p-val<0.05, log2FC>1.5) identified in HEK293 cells and HCT116^*TP53-BirA*/TP53*^, respectively.^39^ Using Cytoscape (version 3.10.3), the complete BioGRID Protein-Protein Interactions network (*H. sapiens*) was imported and the 86 (HEK293) and 78 (HCT116) significantly enriched proteins from each cell line, were used to filter the nodes and edges for specific network construction. A Spatial Analysis of Functional Enrichment (SAFE) was performed for each dataset using an attribute list containing all known GO terms from g:Profiler. A network coverage of 100% and attribute coverage of 92% was reached. SAFE analysis was performed assuming edges of the network are undirected, map-weighted distance, with a 20 percentile threshold. A minimum landscape size of 10 proteins was used to make the composite map with no landscape similarity and a threshold of 0.75. Nodes clustering to an individual domain, were selected and visualized in the SAFE map. For clarity reasons, the remaining proteins were colored light grey. The individual domains were illustrated as a new network and the nodes are colored according to their p-value and log2FC.

## Supplementary figures

**Supplementary figure S1.**
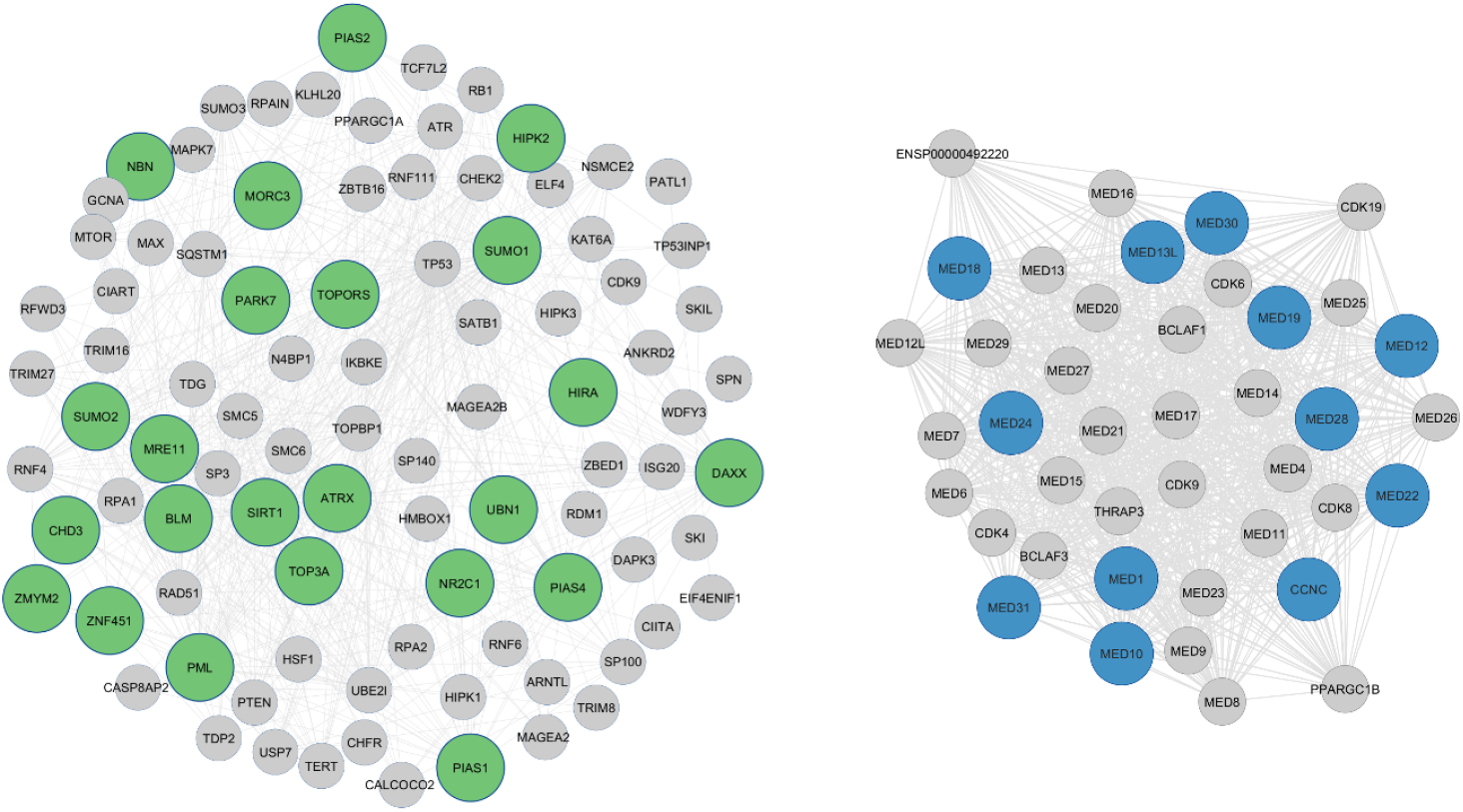
Overview on the enriched proteins from the PML NBs (left) and mediator complex (right) for HEK293 and HCT116, respectively. left panel depicts the 26 enriched proteins found in PML NBs in the HEK293 cell line (FDR=0.01). Right panel illustrates the 12 proteins enriched from the mediator complex for p53 in the HCT116 cell line (FDR=0.01). Complexes were retrieved from the STRING-database and visualized in Cytoscape.

## Author contributions

S.E. conceived and supervised the project. G.V. devised the experimental design, D.D.S. performed the BioID experiments, and E.D.B. performed all analyses and re-analyzed PRIDE data. G.D.M. devised the SAFE analysis for interactomics data used in this work. E.D.B. wrote the original draft of the manuscript. All authors revised the original manuscript.

## Data availability

Proteomics data set is available upon request.

## Competing interests

The authors declare no competing interests.

## Supplementary

**Figure S1.**
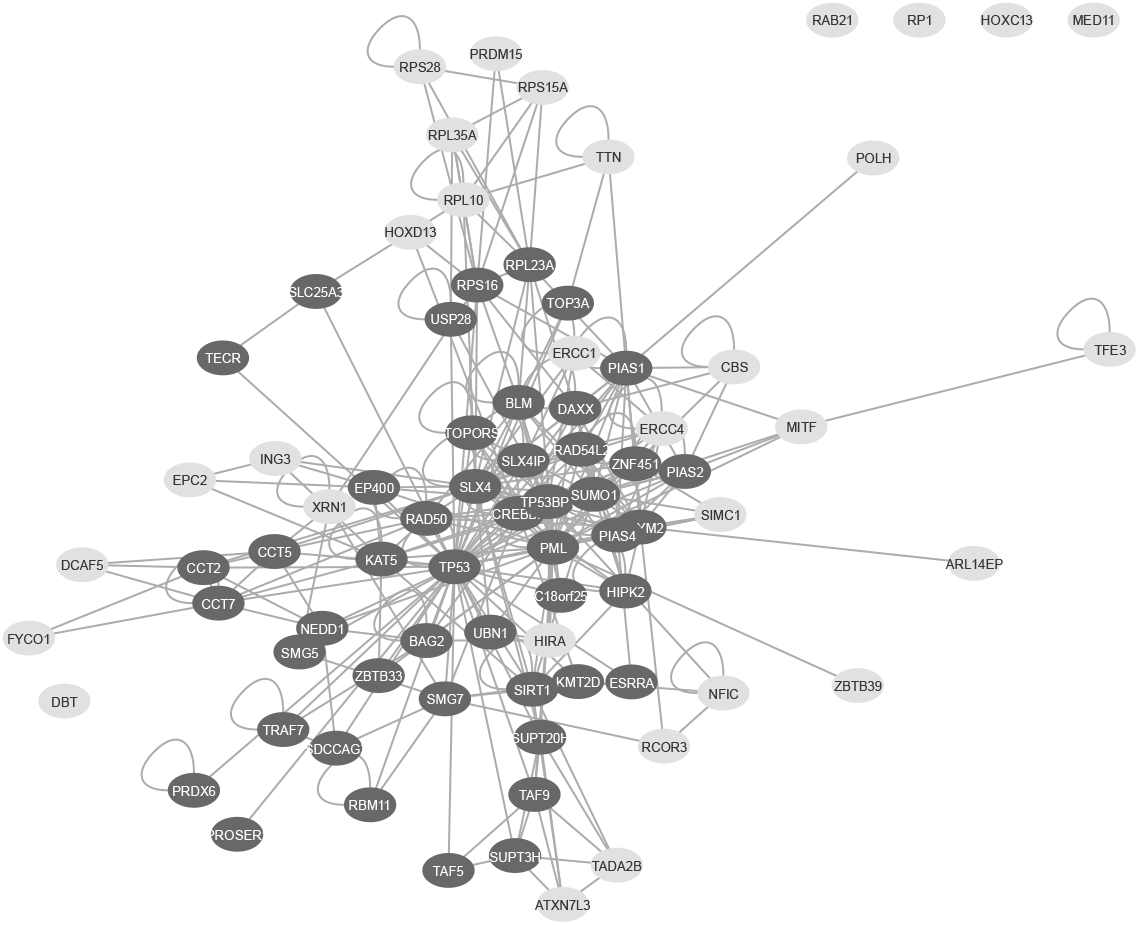
Significantly enriched proteins for p53-BirA*. A total of 86 proteins were identified for p53 compared to control. Out of 86 proteins, 51 (dark grey) have been demonstrated to interact with p53 according to the BioGRID^4.4^ database, while 35 (light grey) have not been reported before.

**Table S1.** Significantly enriched proteins for p53-BirA*. Reported interaction partners of p53 (according to BioGRID^4.4^) have been highlighted.

| *Protein* | *PML nuclear body* | *Histone acetyltransferase complex* | *Transcription coregulator activity* | *SUMO E3 ligases SUMOylate target proteins* |
| --- | --- | --- | --- | --- |
| ARL14EP | No | No | No | No |
| ATXN7L3 | No | Yes | Yes | No |
| BAG2 | No | No | No | No |
| BirA* | No | No | No | No |
| BLM | Yes | No | No | Yes |
| C18orf25 | No | No | No | No |
| CBS | No | No | No | No |
| CCT2 | No | No | No | No |
| CCT5 | No | No | No | No |
| CCT7 | No | No | No | No |
| CREBBP | No | Yes | Yes | Yes |
| DAXX | Yes | No | Yes | Yes |
| DBT | No | No | No | No |
| DCAF5 | No | No | No | No |
| E1B | No | No | No | No |
| EP400 | No | Yes | No | No |
| EPC2 | No | Yes | No | No |
| ERCC1 | No | No | No | No |
| ERCC4 | No | No | No | No |
| ESRRA | No | No | No | No |
| FAM21A | No | No | No | No |
| FDXR | No | No | No | No |
| FYCO1 | No | No | No | No |
| HIPK2 | Yes | No | Yes | Yes |
| HIRA | Yes | No | Yes | No |
| HOXC13 | No | No | No | No |
| HOXD13 | No | No | No | No |
| ING3 | No | Yes | No | No |
| KAT5 | No | Yes | Yes | No |
| KMT2D | No | No | Yes | No |
| MED11 | No | No | Yes | No |
| MITF | No | No | No | Yes |
| NEDD1 | No | No | No | No |
| NFIC | No | No | No | No |
| PIAS1 | Yes | No | Yes | Yes |
| PIAS2 | Yes | No | Yes | No |
| PIAS4 | Yes | No | Yes | Yes |
| PML | Yes | No | Yes | Yes |
| POLH | No | No | No | No |
| PRDM15 | No | No | No | No |
| PRDX6 | No | No | No | No |
| PROSER1 | No | No | No | No |
| RAB21 | No | No | No | No |
| RAB6B | No | No | No | No |
| RAD50 | No | No | No | No |
| RAD54L2 | No | No | No | No |
| RBM11 | No | No | Yes | No |
| RCOR3 | No | No | No | No |
| RP1 | No | No | No | No |
| RPL10 | No | No | No | No |
| RPL23A | No | No | No | No |
| RPL35A | No | No | No | No |
| RPS15A | No | No | No | No |
| RPS16 | No | No | No | No |
| RPS26 | No | No | No | No |
| RPS28 | No | No | No | No |
| SDCCAG3 | No | No | No | No |
| SIMC1 | Yes | No | No | No |
| SIRT1 | Yes | No | Yes | No |
| SLC25A3 | No | No | No | No |
| SLX4 | No | No | No | No |
| SLX4IP | No | No | No | No |
| SMG5 | No | No | No | No |
| SMG7 | No | No | No | No |
| SUMO1 | Yes | No | No | Yes |
| SUMO2 | No | No | No | No |
| SUPT20H | No | Yes | Yes | No |
| SUPT3H | No | Yes | Yes | No |
| TADA2B | No | Yes | Yes | No |
| TAF5 | No | Yes | No | No |
| TAF9 | No | Yes | Yes | No |
| TECR | No | No | No | No |
| TFE3 | No | No | No | No |
| TOP3A | Yes | No | No | No |
| TOPORS | Yes | No | No | Yes |
| TP53 | Yes | No | No | Yes |
| TP53BP1 | No | No | Yes | Yes |
| TRAF7 | No | No | No | No |
| TTN | No | No | No | No |
| UBN1 | Yes | No | No | No |
| USP28 | No | No | No | No |
| XRN1 | No | No | No | No |
| ZBTB33 | No | No | No | No |
| ZBTB39 | No | No | No | No |
| ZMYM2 | Yes | No | No | No |
| ZNF451 | Yes | No | Yes | No |

## Notes

### Competing Interest Statement

The authors have declared no competing interest.

